# Neutrophil to Lymphocyte Ratio (NLR) in captive chimpanzees (*Pan troglodytes*): The effects of sex, age, and rearing

**DOI:** 10.1101/2020.04.27.064717

**Authors:** Sarah J. Neal Webb, Steven J. Schapiro, Chet C. Sherwood, Mary Ann Raghanti, William D. Hopkins

## Abstract

In humans, neutrophil to lymphocyte ratio (NLR) has been used as a clinical tool in diagnosis and/or prognosis of a variety of cancers and medical conditions, as well as in measuring physiological stress over time. Given the close phylogenetic relationship and physical similarities between humans and apes, NLR may similarly be a useful diagnostic tool in assessing chimpanzee health. Only one study has examined NLR in apes, reporting that NLR increased with age and was affected by body-mass index and sex. In the current study, we examined changes in NLR data from longitudinal health records for 443 chimpanzees in two captive chimpanzee populations. Using these data, we analyzed intra-individual changes and inter-individual differences in NLR as a function of age, rearing history, and sex. Contrary to previous studies in humans and the one previous study in chimpanzees, NLR values did not change over a 10-year timespan within individual chimpanzees. However, cross-sectional comparisons revealed a significant quadratic relationship between age and NLR with the highest values during mid-life (20-30 years of age) and the lowest values in younger and older individuals. Additionally, males and mother-reared individuals had higher NLR than females and nursery-reared chimpanzees, respectively. Lastly, males and those with higher NLR values died at younger ages. These findings may suggest that NLR can be used a predictor of longevity in chimpanzees. However, given the complexities of these relationships, more research is needed to determine the utility of NLR as a diagnostic health tool for use in chimpanzees.

## Introduction

Neutrophil to lymphocyte ratio (NLR) is often used as a biomarker of inflammation. The measurement of NLR is obtained through differential cell counts assayed from blood by dividing the number of neutrophils by the number of lymphocytes. Typically, neutrophil counts increase and lymphocyte counts decrease as a function of physiological inflammation, making this ratio a sensitive indicator of inflammation progress [1]. In healthy populations of humans, older individuals have higher NLR, suggesting a possible greater predisposition to inflammation and disease with increasing age [2, 3]. Other studies in healthy humans have found that NLR is moderately heritable (36%) [4], differs as a function of racial disparities in the United States population [2], and that males have higher NLR than females ^[4]^. Average NLR ranges from 1.5 to 2.8 in humans, with reference ranges between 1.1 and 4.5 [1-6].

NLR is often used as a diagnostic and prognostic indicator of a variety of diseases and conditions. NLR can be used to accurately diagnose bacterial sepsis, bacteremia, pancreatitis, and acute appendicitis [7-9]. Higher NLRs are typically associated with poorer outcomes in non-small cell lung disease, acute pancreatitis, pulmonary embolism, cardiac disease, and a variety of cancers, including colorectal, breast, pancreatic, lung, and esophageal, to name a few [10-18]. NLR has also been used to predict all-cause mortality [19, 20], and mortality in patients with specific diseases, including pulmonary embolism [21] and cardiac disease [10]. Elevated NLR may even be connected to Alzheimer’s disease in elderly patients [22, 23].

Apes are physiologically similar to humans, and certain clinical diagnostic and prognostic tools used for humans are also used for chimpanzees [24-26]. Due to recent advances in medical care and captive management, chimpanzee life span has been extended thereby increasing the number of geriatric chimpanzees in captive settings [25, 27]. Like humans, older chimpanzees similarly encounter health conditions and diseases associated with aging including hypertension, multiple types of cancer, heart disease, mobility impairments, arthritis, Alzheimer’s-like pathology, and diabetes [25, 28-31]. Given its utility in humans, NLR may be a valuable measure of health and disease for captive chimpanzees. To date, one study has examined NLR in chimpanzees, finding that NLR was best predicted by age, body mass index (BMI), and an interaction between age, BMI, and sex [32]. Specifically, males had higher NLR than females, but NLR increased more strongly with age and BMI in females than in males. These results are generally consistent with the human literature showing a positive relationship between age and NLR in healthy subjects. However, the sample size in that study was limited to 19 males and 20 females, NLR was sampled at just one point in time for each subject, and the oldest chimpanzee was 31 years old [32]. Since chimpanzees can potentially live up to 60 years in captivity, the relationship of NLR to aging through later stages of the lifespan is unknown.

In addition, captive nonhuman primate health is greatly affected by rearing history. Specifically, nursery-rearing is an early-life stressor that disrupts the development of the immune system, multiple systems in the brain, biobehavioral organization, and HPA axis functioning, among others [33-37]. For example, nursery rearing affects lymphocyte proliferation responses in 2-year old monkeys, which alters immune function and increases susceptibility to infectious diseases later in life [35]. In chimpanzees, the effects of rearing on behavior have been studied extensively, but examinations of the health consequences are sparse. To our knowledge, just one study has examined the relationship between rearing and blood biomarkers of health in chimpanzees [38], finding no differences between mother- and nursery-reared chimpanzees (aged 6 months – 10 years, N = 46) in hematology or serum chemistry values.

Here, we examined relationships between NLR, age, sex, and rearing history in a large, age-diverse sample of captive chimpanzees from two different primate colonies. We calculated NLR at 10 time points (taken once per year during annual physical exams) to examine longitudinal changes in NLR and relationships between NLR, sex, rearing, and age at death. Consistent with previous research in humans and chimpanzees, we predicted that elevated NLR would be exhibited by older individuals, males, and nursery-reared chimpanzees, and that higher NLR values would be associated with younger ages at death.

## Materials and Methods

### Subjects

We collected neutrophil and lymphocyte data from hematology reports for a total of 440 captive chimpanzees (255 females, 185 males) that lived between 1982 and 2019. The NLR data were derived from chimpanzees housed at the National Center for Chimpanzee Care (NCCC, *n* = 204) of the Michale E. Keeling Center for Comparative Medicine and Research at The University of Texas MD Anderson Cancer Center in Bastrop, Texas, and the Yerkes National Primate Research Center (YNPRC, *n* = 236) in Atlanta, Georgia. Chimpanzees ranged in age from 2 to 58 years old (*Mean Age* = 29, *SD* = 12) at the time of their last available data point. All chimpanzees were housed in Primadomes, corrals, or indoor-outdoor runs in social groups. The enclosures contained climbing structures, bedding, and daily environmental enrichment. Care staff fed the chimpanzees a diet of commercially produced primate chow, and fresh fruits and vegetables twice per day. The chimpanzees also had multiple foraging opportunities every day, and ad libitum access to water.

Of the 440 chimpanzees, 182 were mother-reared, 148 were nursery-reared, 102 were wild-born or had an unknown captive rearing history, and 8 had missing data for this variable. For classification purposes, mother-reared individuals were defined as those chimpanzees that were not separated from their mother for at least the first 2.5 years of life and were raised in family social groups of 4 -20 individuals. Nursery-reared chimpanzees were individuals who were separated from the mother within the first month of life due to maternal rejection, illness, or injury. These individuals were cared for by humans, raised in an incubator with access to human infant formula, until they were able to be independent. They were then placed in same-age peer groups until three years of age, at which point they were introduced into larger adult and sub-adult social groups [36, 39]. Those with an unknown rearing history were likely wild-caught, and rearing may have included pet ownership, other methods of human hand-rearing, and/or inclusion in the entertainment industry.

### Neutrophil to Lymphocyte Ratio (NLR)

We used hematology records from annual physical exams (between 1982 and 2019) to obtain values for neutrophils and lymphocytes. Only data from annual physical exams were used for analyses; therefore, any values derived from sedations due to an injury or health issue were not included. However, a small proportion of annual physical exams (approximately 7%) revealed a WBC higher than reported reference ranges for chimpanzees [38], likely indicating a health issue. These WBC counts were not correlated with corresponding NLR values (*p* > 0.10).

Due to missing data for absolute values of neutrophils and lymphocytes, we used percent values to calculate neutrophil to lymphocyte ratios. We used a correlation to examine whether the use of percent values yield different ratios than absolute values, and found that the two methods were positively correlated when using all data points (*r* = .998, *p* < 0.0001, N = 4606), when using a random selection of 20% of the data points (*r* = .995, *p* < 0.0001, N = 925), and when using a random selection of just 5% of the data points (*r* = .999, *p* < 0.0001, N = 232), indicating strong agreement between the two measurements. Therefore, the percent value of neutrophils was divided by the percent value of lymphocytes to obtain the NLR for each chimpanzee at each time point.

Generally, each chimpanzee had one NLR data point per year (corresponding to one annual physical exam per year) across a 10-year period. The 10-year period corresponded to the 10 years prior to the year of the last annual physical exam that preceded either (i) death from natural causes or humane euthanasia, (ii) transfer to another facility, or (iii) the year of 2019.

Where available, we also included NLRs taken at humane euthanasia. We then created 5-year (the **last** five years of NLR data) and 10-year NLR averages for subsequent analyses. These averages did not include any NLR values taken at the point of euthanasia (see below). The age variable used in analyses and shown in figures is the age at each chimpanzee’s last data point (i.e., as stated above, age at the last physical exam, at transfer, or in 2019). Not all 440 chimpanzees had all 5 or 10 years of data. Therefore, the total sample size for 5- and 10-year NLR analyses was 425 and 391 individuals, respectively.

All research and protocol complied with the approved protocols of the UTMDACC and YNPRC Institutional Animal Care and Use Committees, and complied with the legal requirements of the United States and the ethical guidelines put forth by AALAS, the Animal Welfare Act, and The Guide for the Care and Use of Laboratory Animals.

### Data analysis

Histograms and Q-Q plots showed that the data were positively skewed. Exploration of the data revealed five problematic outliers, which were removed from further analyses. We first wanted to examine the effects of age on NLR across the entire sample. Therefore, to examine cross-sectional differences in NLR as a function of age, we used curve estimation to examine both linear and quadratic models for average 5-year and average 10-year NLR. We then used a stepwise regression to determine whether the quadratic function of age (hereafter referred to as quadratic age) explained variance above and beyond the predictors of sex, colony (YNPRC / NCCC), and rearing. To create a dichotomous variable appropriate for a linear regression, and because we were primarily interested in the difference between mother- and nursery-reared individuals, rearing was dummy-coded to compare mother-reared (0) with nursery-reared / wild caught (1); and nursery-reared (0) with mother-reared / wild-caught (1). To examine longitudinal changes in NLR, we used a within-subjects MANCOVA with NLR values from years 1 – 10 as the repeated measure, age (at last physical exam) as the covariate, and sex, colony, and rearing as the between-subject factors (N=362).

Using a subset of the sample for which we had data regarding age at death and NLR values taken at the time of humane euthanasia, we used a linear regression to predict age at death using rearing, sex, and average 5-year NLR (N=180). Lastly, we used paired-samples *t*-tests to examine differences between humane euthanasia NLR values and (i) average 5-year NLR (N=54), and (ii) NLR taken from the last physical exam (N=59). Alpha levels were set at *p* < 0.05 and all analyses were performed using SPSS 26 (IBM Corporation, Chicago, IL, USA). All relevant data are within the manuscript and its Supporting Information files.

## Results

### Cross-sectional analyses

Curve estimation showed significant linear and quadratic relationships between age and average 10-year NLR (linear: *F*[1,389] = 35.43, *p* = 0.0001, *R*^*2*^ = 0.083; quadratic: *F*[2,388] = 26.971, *p* = 0.0001, *R*^*2*^ = 0.122; Fig 1a) and last 5-year average NLR (linear: *F*[1,420] = 28.11, *p* = 0.001, *R*^*2*^ = 0.063; quadratic: *F*[2,419] = 28.82, *p* = 0.0001, *R*^*2*^ = 0.12; Fig 1b). The final model predicting average 10-year NLR with quadratic age, sex, rearing, and colony was significant: *F*(5,387) = 18.14, *p* = 0.0001, *R*^*2*^_*adj*_ = 0.18. The quadratic function of age added uniquely to the model above and beyond other predictors (*R*^*2*^_*change*_ = 0.017, *F*_*change*_[1,382] = 8.15, *p* = 0.005). All predictors were significant (Table 1): (i) Males (mean [*SE*] = 3.06 [0.15]) had higher NLR than females (mean [*SE*] = 2.66 [0.12]; (ii) chimpanzees at the NCCC (mean [*SE*] = 3.04 [0.15]) had higher NLR than those at Yerkes (mean [*SE*] = 2.68 [0.13]); and (iii) mother-reared chimpanzees (mean [*SE*] = 3.34 [0.13]) had higher NLR than nursery-reared chimpanzees (mean [*SE*] = 2.84 [0.13]) and those with an unknown rearing history (mean [*SE*] = 2.40 [0.21]). As shown in Fig 2, mother-reared males had the highest average 10-year NLR values.

**Table 1.** Coefficients in the final models predicting average 10-year NLR and age at death.

|  |  | *b* | *Beta* | *t* | *p* |
| --- | --- | --- | --- | --- | --- |
| Average 10-year NLR | Intercept | 4.578 |  | 10.952 | 0.000 |
|  | Sex | 0.489 | 0.150 | 3.125 | 0.002 |
|  | MR vs. NR&WC | -1.038 | 0.320 | -4.299 | 0.000 |
|  | NR vs. MR&WC | -0.490 | 0.142 | -1.977 | 0.049 |
|  | Colony | -0.361 | -0.112 | -2.272 | 0.024 |
|  | Quadratic Age | 0.000 | -0.176 | -2.854 | 0.005 |
| Age at Death | Intercept | 22.070 |  | 8.952 | 0.000 |
|  | MR vs. NR&WC | 13.592 | 0.557 | 8.463 | 0.000 |
|  | NR vs. MR&WC | 10.241 | 0.403 | 6.416 | 0.000 |
|  | Sex | -5.567 | -0.242 | -4.202 | 0.000 |
|  | 5-year NLR | -0.998 | -.0151 | -2.517 | 0.013 |

Note. MR: Mother-reared; NR: Nursery-reared; WC: Wild-caught.

**Fig 1.**
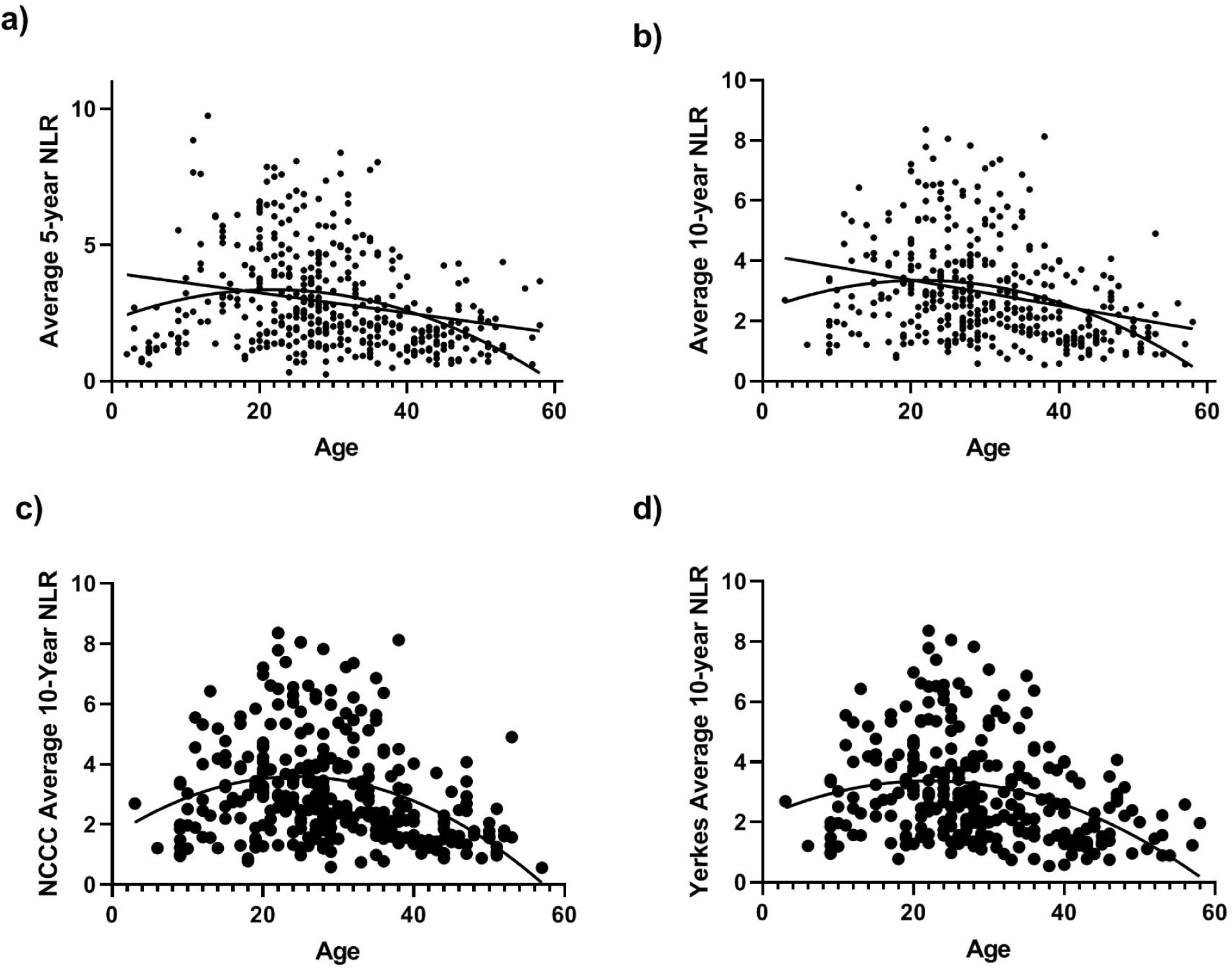
The relationship between chimpanzee age and NLR. Significant linear (*p* = 0.001) and quadratic (*p* = 0.001) relationship between chimpanzee age and average NLR across 10 (a) and 5 (b) years. Quadratic association between age and average 10-year NLR is replicated between NCCC (c) and Yerkes (d) colonies.

**Fig 2.**
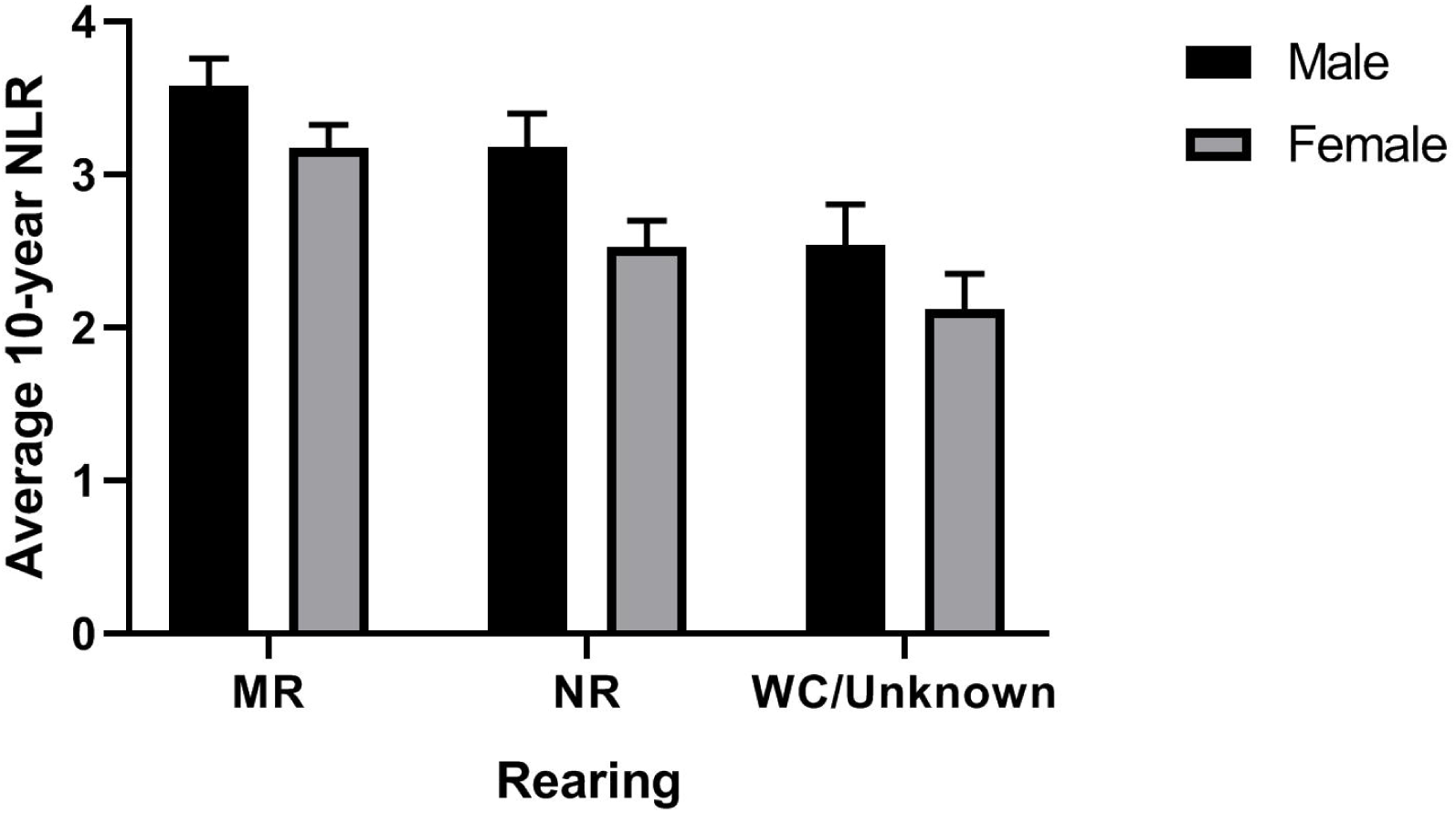
Chimpanzee NLR across rearing types. Average 10-year NLR (+/- SE) as a function of rearing (NR: Nursery-reared; MR: Mother-reared; WC/Unknown: Wild-caught or unknown rearing history) and sex after adjusting for colony and quadratic age.

### Longitudinal analyses

The repeated-measures ANCOVA examining changes in NLR over the course of 10 years while controlling for age showed no change within individuals (*p* = 0.67; Fig 3a-b). Consistent with the regression analysis described above, there was a significant effect of sex, such that males had higher average 10-year NLR values than females (*F*[1,349] = 7.17, *p* = 0.028, Fig 3a). There was also a significant effect of rearing, such that mother-reared chimpanzees had the highest NLR values, followed by nursery-reared, and wild-born chimpanzees (*F*[2,349] = 9.78, *p* = 0.0001, Fig 3b). There were no other significant main or interaction effects.

**Fig 3.**
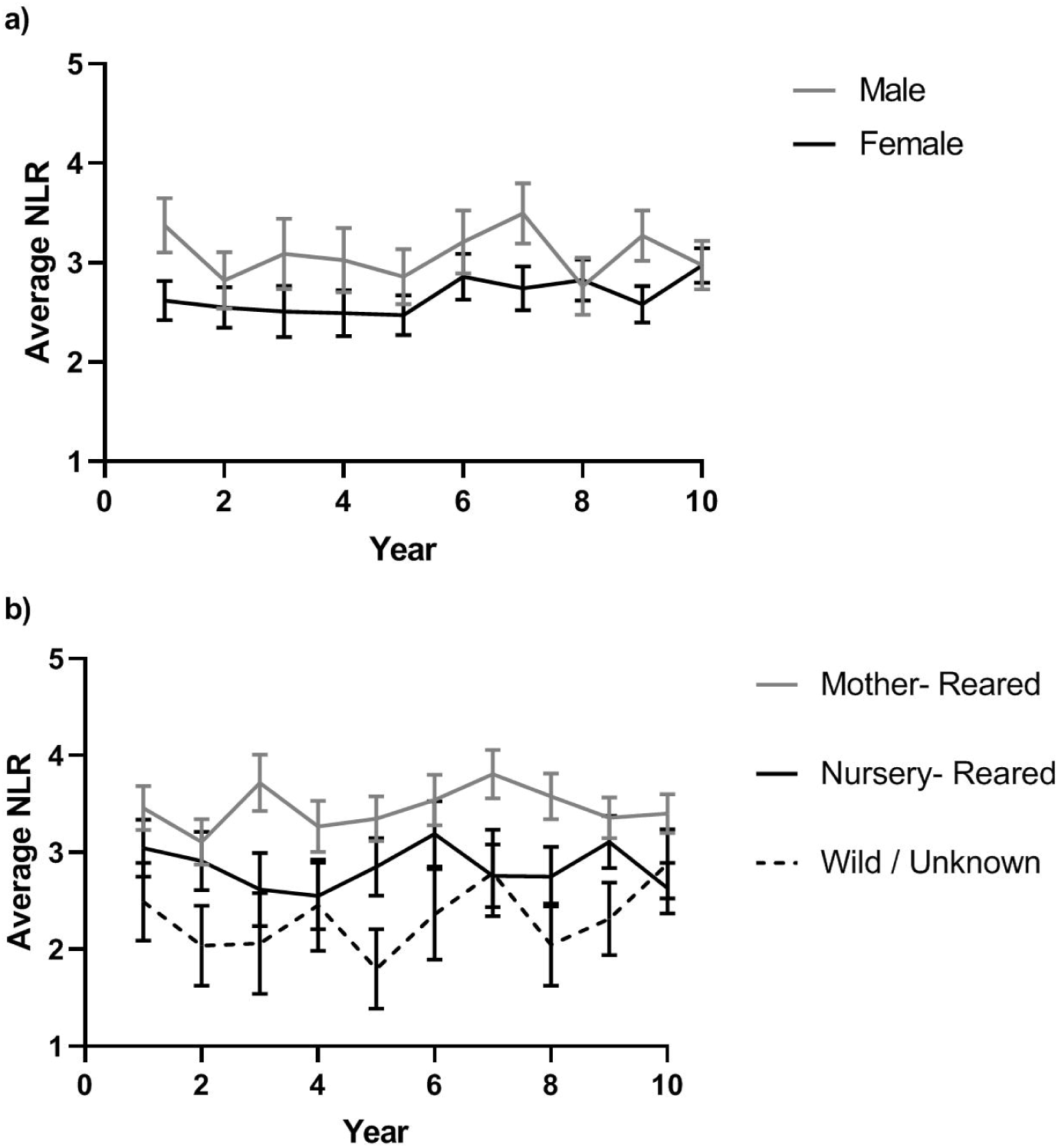
Chimpanzee NLR across sex and rearing. Individual chimpanzee NLR over 10 years as a function of sex (a) and rearing (b). Error bars represent standard error of the mean.

### NLR and mortality

Within chimpanzees for which we had measurements of NLR at the end of their lives, average NLR in the preceding 5-year period and sex were significant predictors of age at death, *F*(4,176) = 35.21, *p* = 0.0001, *R*^*2*^_*Adj*_ = 0.43 (Table 1). As shown in Fig 4a, male chimpanzees and those with higher average NLR values died at younger ages. Additionally, mother-reared individuals (who had the highest NLR values) died at younger ages than nursery-reared individuals, whereas NLR was not related to age at death in wild-caught individuals (not shown in figure). Lastly, NLR at euthanasia was significantly higher (mean [*SE*] = 6.88 [1.23]) than NLR values taken during the last physical exam (mean [*SE*] = 2.28 [0.26]; *t*[58] = 3.61, *p* = 0.001). NLR at euthanasia was also significantly higher than the average NLR for the preceding 5-year period (mean [*SE*] = 2.42 [0.24]; *t*[53)]= 3.49, *p* = 0.001; Fig 4b).

**Fig 4.**
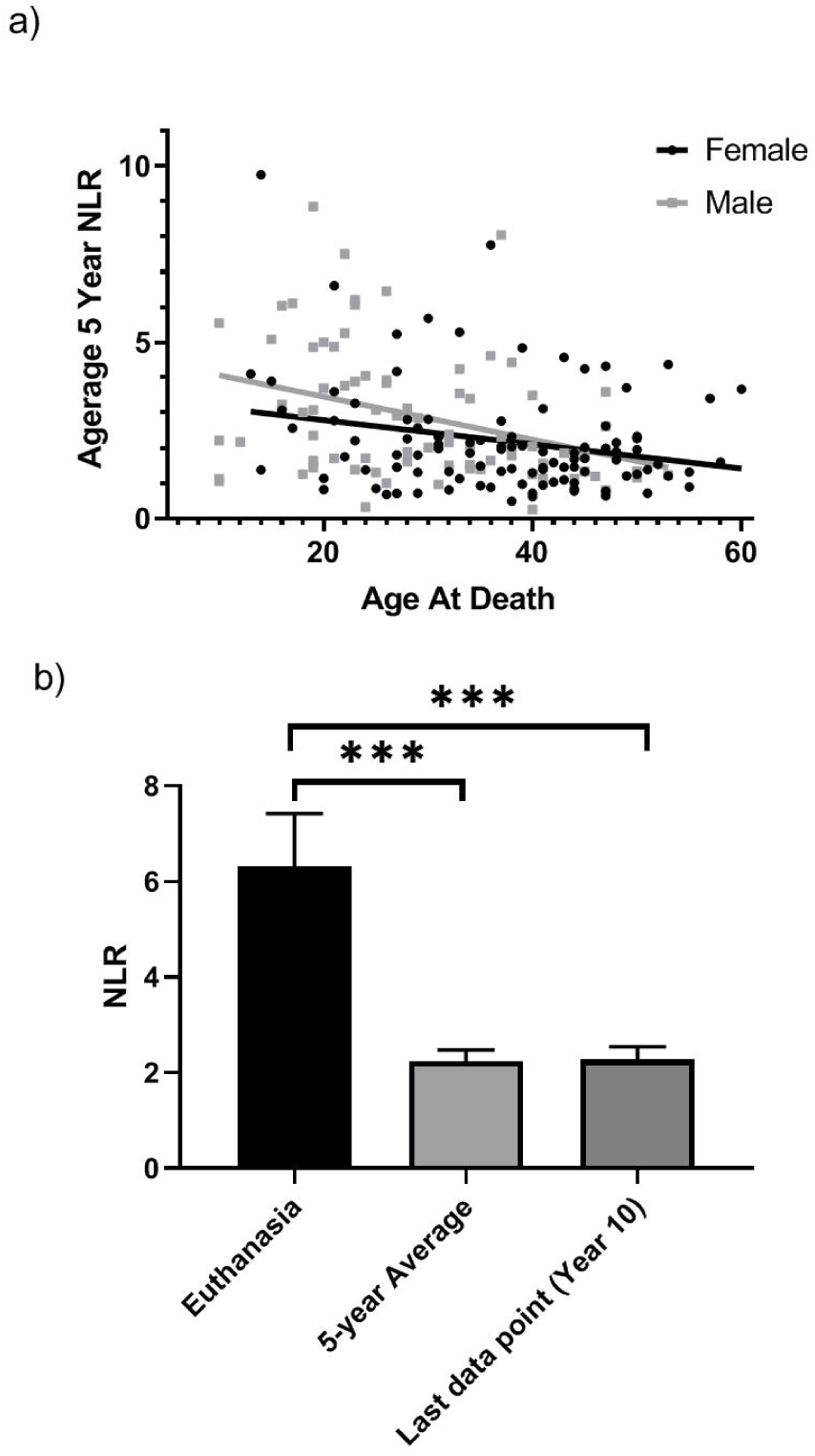
Chimpanzee NLR and mortality. a) Scatterplot showing the negative association between average 5-year NLR and age at death for males and females. b) Mean NLR at euthanasia differs from average 5-year NLR and 10 year NLR (the last point available from a physical exam prior to euthanasia). Error bars represent standard error of the mean. *** *p* = 0.001.

## Discussion

Across a large sample of chimpanzees from two separate colonies, longitudinal analyses showed that, within individuals, NLR did not change over a 10 year period. However, cross-sectional analyses showed that NLR was influenced by sex, rearing, and age. The one study that previously examined NLR in chimpanzees found higher NLR in older individuals (up to 31 years of age) and in chimpanzees with a higher BMI (although older male chimpanzees with a high BMI had lower NLR) [32]. Additionally, males had higher NLR values than females. Results from the current study partly support these results: male chimpanzees also showed higher NLR than females, and NLR was highest in chimpanzees between 25 and 30 years of age. We were able to expand upon these previous findings by using a larger sample size (N = 443 vs. 39), including chimpanzees spanning a larger age range (2 - 58 years old vs. 3 - 31 years), different rearing histories, and using average NLR over several years rather than an NLR value taken at one point in time. In doing so, we found that mother-reared chimpanzees had higher NLR than nursery-reared and wild caught individuals, and there was a significant quadratic relationship between age and average 5- and 10-year NLR in a cross-sectional analysis. We also found that higher average 5-year NLR values predicted younger age at death, and that NLR values at euthanasia were significantly higher than both average 5-year NLR and NLR values at the last physical exam prior to humane euthanasia.

NLR showed a quadratic relationship with age, such that NLR was highest in chimpanzees between 25 and 35 years old, and was lower in both young and old individuals. This is in contrast with some findings in humans, which show that NLR increases linearly with old age in healthy populations, suggesting increased risk for inflammation [1, 3, 6]. The most parsimonious explanation for these data may be that chimpanzees with lower NLRs reach older ages precisely because they maintain better health, and thus, have lower NLR values. Indeed, we found that those with higher NLR values died at a younger age. Perhaps those that have higher average NLRs throughout life [likely indicative of higher levels of inflammation and physiologic stress [40]] die at a younger age, whereas those with lower NLR values live into old age. In this sense, the lower NLR values in elderly individuals may reflect a phenotype associated with “healthy” aging. Interestingly, within individuals, NLR did not increase over a 10-year period, further suggesting that age-associated effects in the cross-sectional sample might be due to subject-specific differences rather than physiological aging. Overall, these results suggest that lower NLR values, which seem to be relatively stable over a 10-year period, may be indicative of longer lifespans in chimpanzees. As such, average 5- and 10-year NLR may serve as a tool aiding in identification of individuals that are at risk for early mortality. Although the NCCC colony had higher average NLRs than the Yerkes colony, the quadratic association between age and NLR was consistent between the two chimpanzee populations (see Fig 1c-d) suggesting that this is a consistent and repeatable finding.

Our data showed that NLR at euthanasia was significantly higher than the NLR data point taken during the preceding physical exam, as well as the average 5-year NLR. These results provide some support for the use of humane euthanasia in captive settings. Specifically, euthanasia is performed for specific, humane reasons, including severe illness or trauma, conditions of chronic wasting, severe cachexia, immobility, organ failure, or moribund state, and upon veterinarian determination that the euthanasia is necessary to alleviate pain and/or distress [41, 42]. This is particularly true for chimpanzees: because of their psychological complexity and phylogenetic proximity to humans, a higher level of ethical and moral justification is required for end-of-life decisions, including assessments of quality of life [41, 42]. The fact that NLR was significantly elevated at the time of euthanasia indicates compromised and/or failing health systems, and thus lends credence to the decision to euthanize under these conditions in which quality of life is low. Indeed, elevated NLR may be used in quality of life programs (similar to the one implemented at the NCCC) to help identify which animals may be closer to their endpoints prior to the need for euthanasia.

Consistent with previous research showing the utility of NLR as a diagnostic tool in humans, NLR can be used as a tool to aid in diagnosing severe illness and trauma. In combination with a multitude of diagnostic criteria used to identify clinical illnesses, the utility of NLR may be in its use as a prognostic indicator of shorter lifespan and in identifying at-risk individuals. For example, the leading cause of death in chimpanzees is cardiac disease, and research has shown that males are more likely to suffer from myocardial fibrosis and related sudden death [24, 43-47]. Furthermore, many of these sudden deaths occur mid-life, between 25-30 years of age [44, 45, 47]. The finding that NLR was higher in males, and was highest in individuals between 25 and 35 years of age seems consistent with this cardiac death, sex, and age pattern. Interestingly, we also found that mother-reared chimpanzees had higher NLR values and died at younger ages than nursery-reared chimpanzees (as well as wild-caught chimpanzees, but that effect is likely due to the confounding factor that wild-caught chimpanzees are significantly older than chimpanzees of other rearing types). This finding is surprising: given the multitude of health consequences associated with nursery-rearing, we expected to find that nursery-reared individuals would have the highest NLR. Overall, if NLR indeed indicates increased inflammation and disease risk, this would suggest that mother-reared individuals, particularly mother-reared males between 25 and 35 years of age, have the highest risk for negative outcomes. Unfortunately, we are currently unable to speculate about the mechanisms underlying this increased risk. Regardless, given the similarity between the patterns described above, NLR may be prove a useful biomarker in identifying individuals at risk for sudden cardiac death.

Though NLR values differed as a function of age, sex, rearing, and colony, collectively these factors explained only 18% of the variation in NLR. As such, there are likely a multitude of other factors, both genetic and environmental, affecting NLR. Because previous studies have found that NLR in humans is moderately heritable [4], we are currently examining the heritability of NLR in chimpanzees. Additionally, it is possible that individual differences in genetic or biological aging mechanisms may help to explain individual differences in NLR. For example, there is significant variation between the biological (as measured through changes in epigenetic methylation) and chronological age of chimpanzees [48]. Animals that show a faster rate of biological aging may have shorter lifespans. Additionally, other measures or indicators of physiological chronic stress are likely correlated with NLR. For example, allostatic load, a measure of stress-induced physiological damage over the lifetime, is higher in wild-caught gorillas compared to their mother- and nursery-reared counterparts [49, 50]. Given the lower NLR values of wild-caught chimpanzees in the current study (although, again, this may be confounded with age), it is possible that chimpanzees with lower allostatic load also have lower NLR. Additional research examining these variables would shed light on healthy aging in chimpanzees, a topic that is of increasing importance given the longer lifespans and aging populations of captive chimpanzees [25, 30]. Regarding environmental factors, additional studies are currently underway that aim to examine the effects of chronic conditions, past experimental history, and body condition scores (a proxy measure of BMI) on NLR values.

In summary, although these findings reveal a complex relationship between NLR and individual chimpanzee characteristics, age, rearing, and sex explain only a modest amount of the total variance in average 10-year NLR values. The current study builds on previous findings in chimpanzees by showing that (i) the oldest chimpanzees (up to 58 years of age) had lower NLR values, (ii) mother-reared males had the highest NLR values, and (iii) individuals with higher NLR values had shorter lifespans. We believe that these older chimpanzees have longer lifespans precisely because of their lower NLR values; however, why some individuals have lower NLR values than others is a research question that we hope to continue exploring. Much more research is needed to understand the genetic and environmental factors that affect NLR, the relationships between NLR, lifespan, and clinical illness, as well as the use of NLR as a diagnostic and prognostic tool. Additionally, although the current study provides a preliminary reference for normal male and female NLR values [3.10 and 2.58, respectively, consistent with previous research: mean NLR values of 2.66 and 2.54 [32], respectively], more research is needed to confirm the range of normal NLRs in chimpanzees. This will allow identification of atypical and/or clinically abnormal values that may signify problems and/or warrant intervention.

## Acknowledgements

The authors would like to thank Mary Ann Cree for assistance in obtaining archival Yerkes data, as well as Dr. Michele Mulholland and Mary Catherine Mareno for helpful comments during preparation of this paper. We would also like to thank the care staff at the National Center for Chimpanzee Care and the Yerkes National Primate Research Center for their unwavering care of the chimpanzees.

